# (*R*)-*S*-Adenosyl-L-methionine hydrolases counter sulfonium epimerisation in thermophilic archaea

**DOI:** 10.1101/2025.08.29.672840

**Authors:** A. Bartels, M. K. F. Mohr, P. Nußbaum, B. Wassmer, L. Rasquin, S. V. Albers, J. N. Andexer

**Affiliations:** University of Freiburg, Institute of Pharmaceutical Sciences, Albertstr. 25, 79104 Freiburg (Germany); University of Freiburg, Faculty of Biology, Molecular Biology of Archaea, Schänzlestr. 1, 79104 Freiburg (Germany)

**Keywords:** SAM hydrolase, epimerisation, thermophiles, DUF62, stereoselectivity

## Abstract

S*-Adenosyl-L-methionine (SAM) is the second most used enzyme cofactor and vital for numerous cellular reactions such as methylation or polyamine synthesis. While most stereocentres of the biologically active (*S_S_,S_Cα_*)-SAM are fixed, epimerisation at the methyl sulfonium centre is driven by heat, yielding biologically inactive (*R_S_,S_Cα_*)-SAM. This SAM diastereomer disturbs SAM-dependent pathways, posing a metabolic threat especially to thermophilic organisms*. In vitro *analysis shows that SAM hydrolases cleave the biologically inactive (*R_S_,S_Cα_*)-SAM, thereby constituting to a metabolic salvage pathway. For further analysis of the biological relevance, we characterised two archaeal SAM hydrolases from the thermophilic* Sulfolobus acidocaldarius *and the halophilic* Haloferax volcanii, *confirming their selectivity towards (*R_S_,S_Cα_*)-SAM* in vitro. *Genetic manipulation in the native hosts supports a significant role of the SAM-hydrolases in decreasing the share of intracellular (*R_S_,S_Cα_*)-SAM to sustain cellular functions in thermophilic organisms*.

## Introduction

*S*-Adenosyl-L-methionine (SAM) is one of the most used enzyme cofactors and is primarily known as a source of methyl groups. It also is a precursor for polyamine synthesis, and a cofactor for radical SAM enzymes sustaining vital cellular reactions throughout all three kingdoms of life.^[1]^ SAM contains six stereocentres, four of which are on the carbon atoms of the ribosyl moiety. The remaining two are located at the α-carbon (*C*_α_) of the amino acid moiety and at the sulfonium (*S*) group which is essential for all mechanisms of SAM-dependent enzymes. The five carbon stereocentres of SAM, both at the amino acid moiety and the adenosyl moiety, are fixed. However, pyramidal inversion at the sulfonium enables the cofactor to spontaneously epimerise to (*R*_S_,*S*_Cα_)-SAM (**Figure 1**).^[2–4]^ While many studies on SAM-dependent enzymes show the use of the biologically diastereomer, active the (*S*_S_,*S*_Cα_)-SAM epimerised (*R*_S_,*S*_Cα_)-SAM diastereomer was reported to be biologically inactive for methyl transferases (MTs) and to even inhibit some SAM-dependent enzymes *in vitro*.^[3]^ Effects of accumulating (*R*_S_,*S*_Cα_)-SAM on whole organisms where not investigated thoroughly so far. Under physiological conditions, epimerisation of SAM is slow, however, higher temperatures promote inversion at the sulfonium, which makes especially thermophiles prone for elevated (*R*_S_,*S*_Cα_)-SAM concentrations.^[3–5]^

**Figure 1.**
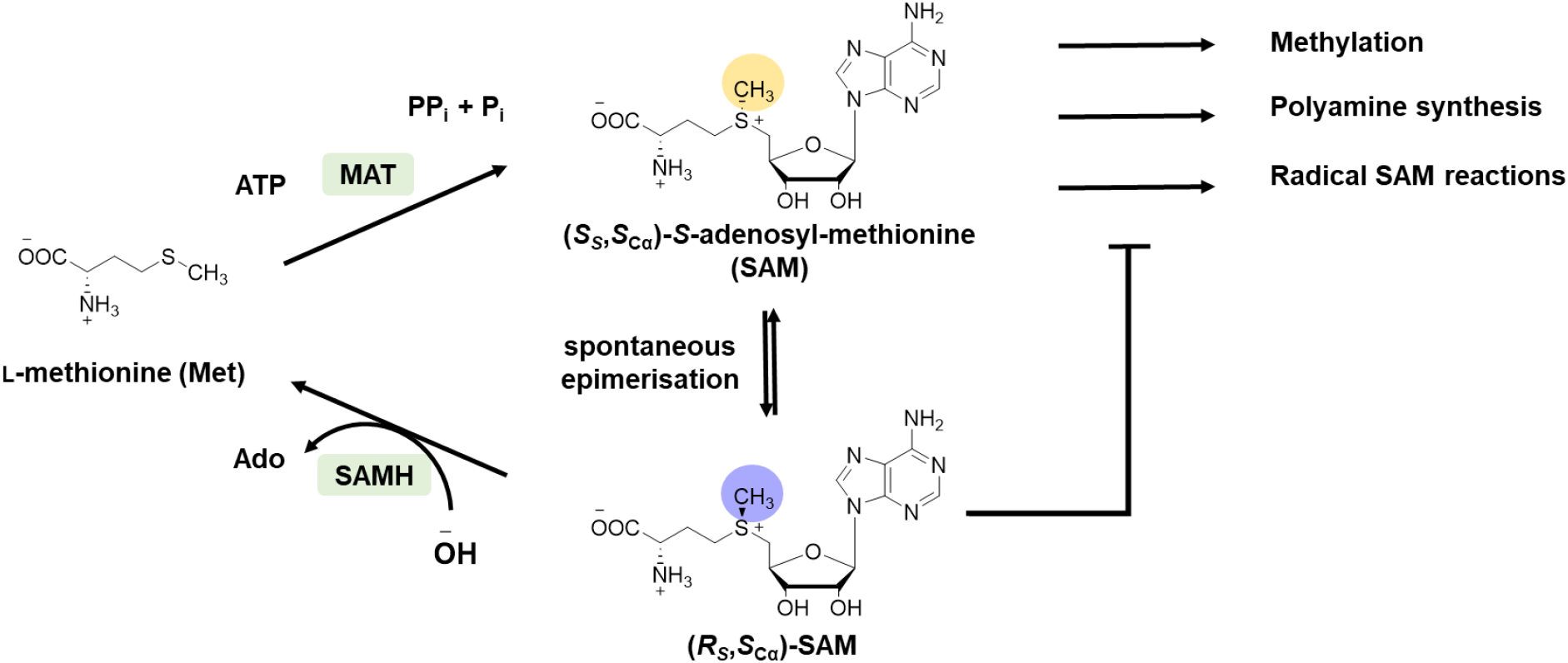
Synthesis and postulated degradation of SAM. SAM sulfonium-diastereomers and SAMH reaction in the context of cellular transmethylation reactions.

In addition to epimerisation, SAM degrades under physiological conditions by two additional non-enzymatic mechanisms: Intramolecular attack of the carboxy group at the C_γ_ of the amino acid moiety leads to formation of homoserine lactone and 5′-methylthioadenosine (MTA). Alternatively, deprotonation at C5′ through water and subsequent cleavage of the glycosidic bond yields *S*-ribosyl-L-methionine and adenine.^[4,6]^

In order to cope with the fast degradation of the cofactor, SAM is rapidly consumed after its synthesis in nature. Methionine adenosyltransferases (MATs) stereo-specifically synthesise (*S*_S_,*S*_Cα_)-SAM from adenosine 5′-triphosphate (ATP) and methionine (Met).^[7]^ It has been shown that the supply of Met can increase the production of SAM in different microorganisms.^[8–11]^ While the cofactor is rapidly used after its production, small shares of (*R*_S_,*S*_Cα_)-SAM have been found in cellular extracts, e.g. 3% in mouse liver extract.^[6,12]^ One of nature’s options to lower (*R*_S_,*S*_Cα_)-SAM concentrations is SAM hydrolases (SAMHs), catalysing the hydrolysis of SAM to Met and adenosine (Ado) (**Figure 1**).^[13]^ *In vitro* studies on the SAMH from the marine bacterium *Salinispora tropica* showed the stereoselective degradation of (*R*_S_,*S*_Cα_)-SAM from an epimerised mixture of the cofactor. This suggests that SAMHs constitute a salvage pathway to recover the metabolites from the (*R*_S_,*S*_Cα_)-SAM diastereomer and prevent inhibition of the organism’s metabolism through the epimerised cofactor.^[14–17]^ A second known (*R*_s_,*S*_Cα_)-SAM salvage pathway is constituted by homocysteine *S-*methyltransferases (HSMTs) which synthesise methionine from homocysteine and (*R*_s_,*S*_Cα_)-SAM.^[18,19]^ Elevated levels of (*R*_S_,*S*_Cα_)-SAM in SAM salvage pathway deficient organisms and a physiological role of SAMHs to prevent intracellular (*R*_S_,*S*_Cα_)-SAM accumulation still have to be confirmed.

Epimerisation of SAM strongly increases with an increase in temperature. Therefore, we were focusing on the investigation of the role of SAMH within organisms growing at varying temperatures. Archaea are microorganisms that constitute one of the three domains of life. They are classified in four superphyla and often (while not exclusively) inhabit extreme environments making them interesting for biotechnological application and investigating cellular functions.^[20,21]^ The availability of archaeal model organisms with adequate genetic tools, growing at moderate and high temperatures, as well as the strong conservation of SAMHs in archaea suggests them to be optimal hosts for investigating the temperature-dependent role of SAMHs in the maintenance of intracellular (*R*_S_,*S*_Cα_)-SAM concentrations.

For our investigation, we selected the thermoacidophilic crenarchaeon *Sulfolobus acidocaldarius*, thriving at a temperature of 75 °C, and the halophilic euryarchaeon *Haloferax volcanii*, which has an optimal growth temperature of 45 °C.^[21,22]^ First, we recombinantly produced and purified SAMHs from *S. acidocaldarius DSM639 and H. volcanii H26* in *Escherichia coli* BL21 Gold and analysed the stereoselectivity of the SAM hydrolysis *in vitro*. To investigate their role in maintaining (*R*_S_,*S*_Cα_)-SAM level, we deleted the genes encoding SAMHs in both organisms and analysed the influence on the intracellular SAM content as well as the respective share of (*R*_S_,*S*_Cα_)-SAM.

## Results

### *In vitro* activity of *Saci*SAMH and *Hv*SAMH

The gene, *saci0719*, encoding *Saci*SAMH was identified as part of a conserved operon (*saci0718* – *saci0720*) containing a triphosphate tunnel metalloenzyme and a nicotinamide mononucleotide adenylyltransferase, located close to one origin of replication.^[23]^ The SAMH of *H. volcanii* (*hvo_0781*) was identified as a homologue of *saci0719 via* sequence similarity search using NCBI BLAST. The identified genes were cloned into modified pET15b vectors carrying the sequence for a His_6_-tag (*Saci*SAMH) or a Strep-tag (*Hv*SAMH), recombinantly expressed in *Escherichia coli* BL21 Gold and the proteins obtained were purified *via* the corresponding affinity chromatography. The purified proteins were then analysed towards their catalytic activity to cleave (*R*_S_,*S*_Cα_)-SAM from an epimerised SAM mixture *in vitro*. Due to the instability of SAM at physiological pH, assays were performed at 37°C and pH 7.5 for all SAMHs. Both, *Saci*SAMH and *Hv*SAMH showed activity towards the cleavage of (*R*_S_,*S*_Cα_)-SAM to Ado as observed by HPLC-UV (**Figure 2**). Reactions with *Hv*SAMH showed the formation of a side product (**Figure 2 B**). The retention time suggests it to be the deamination product inosine (Ino), the molecular basis for this is currently investigated in our laboratory.

**Figure 2.**
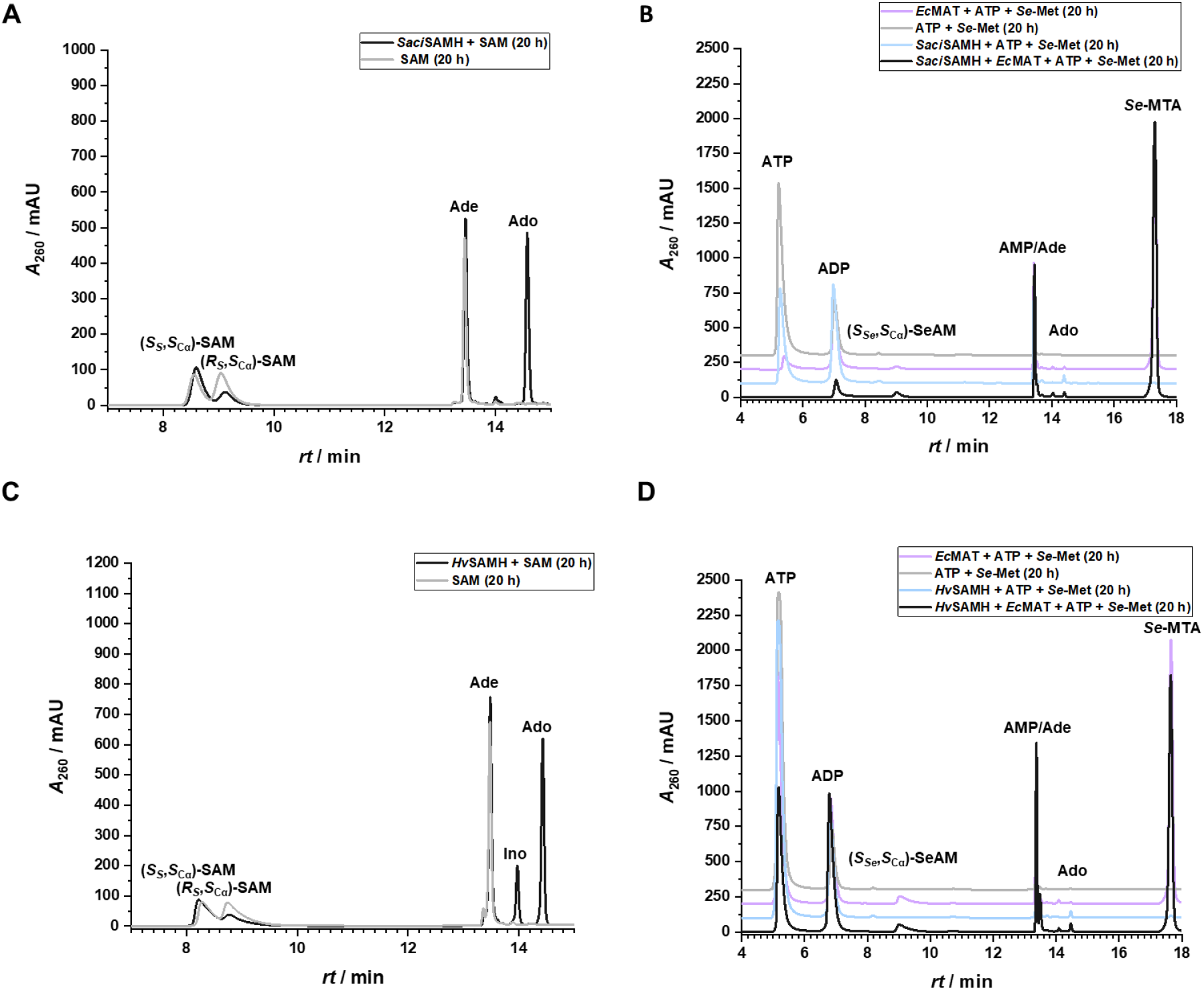
Stereoselective cleavage of (*R*_S_,*S*_Cα_)-SAM by archaeal SAMHs. HPLC-UV chromatograms at *λ* = 260 nm showing results of *in vitro* assays at 37 °C after 20 h. Ado formation from epimeric mixture of SAM is observed for *Saci*SAMH (**A**) and *Hv*SAMH (**C**), only (*R*_S_,*S*_Cα_)-SAM is cleaved. Cascade reactions with *Ec*MAT show the formation of (*S*_Se_,*S*_Cα_)-SeAM from *Se*-Met and ATP, but no Ado formation for either *Saci*SAMH (**B**) or *Hv*SAMH (**D**) confirming the SAMHs’ specificity for the (*R*,*S*)-diastereomer of the cofactor. ADP: adenosine 5′-diphosphate, AMP: adenosine 5′-monophosphate. Offset between chromatograms in B and D is 100 mAU.

For confirmation of the stereospecificity of the reaction, the SAMHs were tested in a cascade starting with the MAT of *E. coli* (*Ec*MAT), ATP and L-selenomethionine (instead of ATP and L-methionine). Selenium, which is less electronegative compared to sulfur, leads to a higher pyramidal inversion barrier and, hence, a SAM analogue with stable configuration at the chalcogen-onium.^[5]^ This allows for the analysis of the SAMHs’ ability to cleave (*S*_Se_,*S*_Cα_)-SeAM. In the respective assays, Ado formation was observed for neither *Hv*SAMH nor *Saci*SAMH (**Figure 2 C** and **D**). The successful (*S*)-*Se-*adenosyl-L-selenomethionine (*Se*AM) production by *Ec*MAT was confirmed by the corresponding peak in the HPLC chromatogram. Based on literature showing many SAM-dependent enzymes to also convert SeAM, it can be assumed that the lack of Ado formation from (*S_Se_*,*S_Cα_*)-SeAM is caused by the configuration of the cofactor rather than the sulfonium to selenium substitution.^[24,25]^ Therefore, the same stereospecificity for the (*R*_Se_,*S*_Cα_)-configuration can be inferred for the sulfur-containing cofactor.

SAM-dependent chlorinases are close relatives to the SAMHs, which catalyse a nucleophilic attack of chloride ions on 5′-C of (*S*_S_,*S*_Cα_)-SAM, producing 5′-chloro-5′-deoxyadenosine (5CldA). Despite the high concentration of chloride ions, 5CldA was not produced by any of the SAMHs, resulting in a specific activity towards the cleavage of (*R*_S_,*S*_Cα_)-SAM (**Figure S3**).

### Biological role of SAMHs

Having confirmed stereospecific cleavage of (*R*_S_,*S*_Cα_)-SAM by the archaeal SAMHs *in vitro*, we further investigated their biological role. Deletion strains for the respective SAMHs were designed for *S. acidocaldarius* (*Saci_KO*) and *H. volcanii* (*Hv_KO*). To our surprise, the KO strains showed no obvious change in morphology or viability under standard culturing conditions. For further analysis, the influence of SAMHs on the intracellular SAM content of the archaea was investigated. The focus of these experiments was to determine the share of (*R*_S_,*S*_Cα_)-SAM in the total SAM content.

The share of (*R*_S_,*S*_Cα_)-SAM was higher in *S. acidocaldarius* (WT: 8%) compared to *H. volcanii* (WT: not detectable) (**Figure 3 A** and **B**). This is in line with an expected faster epimerisation rate of SAM at elevated temperatures.^[4]^ When the *samh* genes were deleted in the KO strains, the share of (*R*_S_,*S*_Cα_)-SAM increased significantly to 17% for *Saci_KO* and to around 5% for *Hv_KO*, highlighting the threat of (*R*_S_,*S*_Cα_)-SAM accumulation and confirming the intracellular activity of the SAMHs in the maintenance of a low (*R*_S_,*S*_Cα_)-SAM level (**Figure 3 A** and **C**).

**Figure 3.**
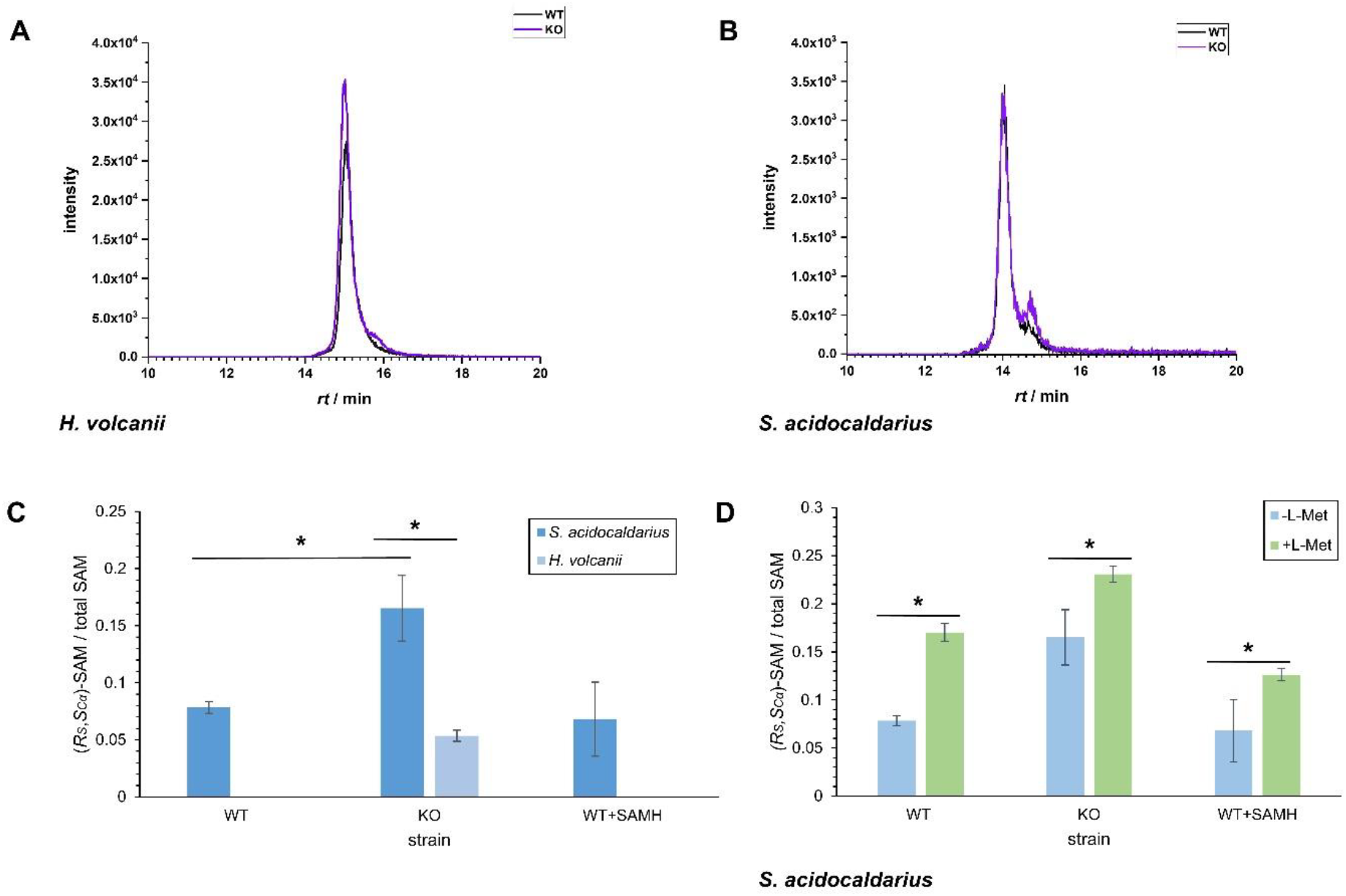
Intracellular SAM content in WT and KO strains. **A**) Exemplary extracted ion chromatograms of MRM-scan (*m/z* 399.15 / *m/z* 250.02) on samples of *H. volcanii*. **B**) Exemplary extracted ion chromatograms of MRM-scan (*m/z* 399.15 / *m/z* 250.02) on samples of *S. acidocaldarius*. **C**) Share of (*R*s,*S*_Cα_)-SAM in total intracellular SAM content of *S. acidocaldarius* as detected via LC-MS/MS (*m/z* 399.15 / *m/z* 250.02). Values are means of biological triplicates; SD is shown as error bars.* = p < 0.05 in two-tailed unpaired t-test.**D**) Influence of L-Met addition on share of (*R*s,*S*_Cα_)-SAM in total intracellular SAM content of *S. acidocaldarius* as detected via LC-MS/MS (*m/z* 399.15 / *m/z* 250.02). Values are means of biological triplicates; SD is shown as error bars. * = p < 0.05 in two-tailed unpaired t-test.

To further push the accumulation of (*R*_S_,*S*_Cα_)-SAM, we supplied exponentially growing *S. acidocaldarius* cultures with 10 mM Met which was reported to increase intracellular SAM concentrations in *E. coli*.^[8–10]^ As predicted, the total SAM level increased for all cultures, indicating that Met is one limiting factor for SAM synthesis also in archaea. Supply of Met increased the share of (*R*_S_,*S*_Cα_)-SAM significantly to 17% for *Saci_WT* and a maximum of 23% for *Saci_KO* (**Figure 3 D**). The elevated share of (*R*_S_,*S*_Cα_)-SAM could be reduced to 13% by overexpression of *saci_samh* in *Saci_WT*. These data suggest an important role of SAMHs for cellular maintenance of (*R*_S_,*S*_Cα_)-SAM concentrations, especially at high temperatures. They also show the potential of *Saci_WT* in studying the effects of (*R*_S_,*S*_Cα_)-SAM accumulation on SAM-dependent metabolic pathways and investigating strategies to counter accumulated (*R*_S_,*S*_Cα_)-SAM by *samh* overexpression.

## Discussion and Outlook

With these data, we verified that the identified enzymes of the DUF62 superfamily from *S. acidocaldarius* and *H. volcanii* are (*R*_S_,*S*_Cα_)-specific SAMHs. This stereoselectivity falls in line with the results of KORNFUEHRER *et al*. for the homologous enzyme from *Salinispora tropica* and strengthens the hypothesis that SAMHs can constitute an (*R*_S_,*S*_Cα_)-SAM salvage pathway.

Deletion of the *samh* genes in *S. acidocaldarius* and *H. volcanii* did not directly influence the morphology or growth of the strains. However, the share of (*R*_S_,*S*_Cα_)-SAM strongly increased when the *samh* genes were deleted supporting the hypothesis of SAMHs to be involved in salvage of the biologically inactive diastereomer. Under standard conditions, we could not observe effects of accumulating (*R*_S_,*S*_Cα_)-SAM on the physiological appearance of the KO strains, however, effects on SAM-dependent pathways such as general methylation activity still need to be analysed. As SAM is involved in numerous metabolic and regulatory pathways, a change in behaviour under stress (e.g. elevated temperature, oxidative stress, starvation, UV-radiation) is very likely. For confirmation of these hypotheses, the investigation of SAMHs in further organisms and under various culturing conditions is needed.

The share of (*R*_S_,*S*_Cα_)-SAM in *S. acidocaldarius* lies with 8% for the WT strain more than two-times above the values previously found in mouse liver extracts (3%) and rat brain (3%), supporting the report of higher SAM epimerisation at elevated temperatures.^[6,17]^ Thus, we added to the hypothesis that (*R*_S_,*S*_Cα_)-SAM accumulates more easily within organisms living at elevated temperatures (17% (*R*_S_,*S*_Cα_)-SAM in *Saci_KO vs*. 5% (*R*_S_,*S*_Cα_)-SAM in *Hv_KO*). By the supply of Met and overexpression of *samh*, different levels of (*R*_S_,*S*_Cα_)-SAM accumulation could be generated. Using these models, we will further investigate the influence of (*R*_S_,*S*_Cα_)-SAM accumulation on crucial SAM-dependent pathways in archaea.

## Materials and Methods

All chemicals were purchased from Sigma Aldrich, Carl Roth or Thermo Scientific, except stated otherwise.

### Media preparation for *H. volcanii and S. acidocaldarius*

#### H. volcanii

Stock solutions were prepared as follows. For 2 L of a 30% buffered Salt Water (BSW) stock solution 480 g NaCl, 60 g MgCl_2_ x 6 H_2_O, 70 g MgSO_4_ x 7 H_2_O, 14 g KCl were first dissolved in ∼1.6 L warm distilled water using a magnetic stirrer. Then 40 mL of 1 M Tris-HCl pH 7.5 was added and topped up with distilled water to a final volume of 2 L.

10x YPC stock solution was prepared by dissolving 8.5 g yeast extract (Difco), 1.7 g Peptone (Oxoid) and 1.7 g of Casamino acids (Oxoid) in 170 ml distilled water. The pH of the stock was adjusted to 7.2 by addition of KOH and subsequently autoclaved.10x CA stock solution was prepared by dissolving 10 g of Casamino Acids (Oxoid) to ∼150 mL of distilled water. Then the pH was adjusted to 7.2 by addition of 1 M KOH. Distilled water was then added to a final volume of 200 mL and the stock solution autoclaved. 1000x vitamin solution was prepared by dissolving 50 mg Thiamine and 5 mg Biotin in 50 mL of distilled water. The solution was filter sterilized. 100x trace element solution was prepared by dissolving 5 g EDTA (disodium salt), 0.8 g FeCl_3_ (4.9 mmol), 0.05 g ZnCl_2_ (0.37 mmol), 0.01 g CuCl_2_ (0.074 mmol), 0.01 g CoCl_2_ (0.077 mmol), 0.01 g H_3_BO_3_ (0.16 mmol), 1.6 g MnCl_2_ (12.7 mmol), 0.01g NiSO_4_ (0.065 mmol), 0.01 g H_2_MoO_4_ (0.062mmol) in 1L of distilled water.^[26]^

Once everything was dissolved the pH was adjusted to 7 and the solution filter sterilized.

To prepare medium, 200 mL of 30% BSW, 100 mL of distilled water were mixed and autoclaved. For selective CAB-medium, 33 mL of 10x CA stock, 2 mL of a 0.5 M CaCl2, 3.3 mL of 100x trace-element solution and 0.33 mL of 1000x Vitamin solution were added. For non-selective YPC-medium 33 mL of 10x YPC stock and 2 mL of 0.5 M CaCl2 were added.

Plates for *H. volcanii* were prepared as described before.^[27]^

#### S. acidocaldarius

Stock solutions were prepared as follows. For 2 L of a 10× Brock basal salts stock solution, 26 g (NH_4_)_2_SO_4_, 5.6 g KH_2_PO_4_, and 5 g MgSO_4_·7H_2_O were dissolved in ∼1.6 L of distilled water using a magnetic stirrer (these amounts correspond to 1.3 g/L, 0.28 g/L, and 0.25 g/L in the final 1× medium). The solution was then brought up to 2 L with water and autoclaved. A 10× N-Z-Amine stock solution was prepared by dissolving 10 g of N-Z-amine (casein hydrolysate) in ∼150 mL of distilled water. The pH of this stock was adjusted to ∼7.0 with KOH, then distilled water was added to a final volume of 200 mL and the solution autoclaved. A 10× dextrin stock solution was prepared by dissolving 20 g of dextrin in ∼300 mL of distilled water, adjusting the pH to ∼7.0 with KOH, and adding water to a final volume of 400 mL before autoclaving. A 100× trace element solution was prepared by dissolving 0.2 g FeCl_3_·6H_2_O, 0.2 g MnCl_2_·4H_2_O, 0.5 g Na_2_B_4_O_7_·10H_2_O, 22 mg ZnSO_4_·7H_2_O, 5 mg CuCl_2_·2H_2_O, 10 mg NiSO_4_·6H_2_O, 3 mg Na2MoO4·2H_2_O, 3 mg VOSO_4_·xH_2_O, and 1 mg CoSO_4_·7H_2_O in 1 L of distilled water (these trace components are based on the modified Brock medium). A few drops of concentrated H_2_SO_4_ were added to the trace solution to bring the pH to ∼1–2 (to keep iron and other metals in solution), and the solution was filter-sterilized (0.2 µm). Additionally, a 0.5 M CaCl_2_ stock solution was prepared separately and sterilized by autoclaving.

To prepare 1 L of **Brock medium** for *S. acidocaldarius*, 100 mL of the 10× basal salts stock and ∼600–700 mL of distilled water were mixed in a glass bottle and autoclaved. After cooling, 100 mL of the 10× N-Z-Amine stock, 100 mL of the 10× dextrin stock, 10 mL of the 100× trace element solution, and 1 mL of the 0.5 M CaCl2 stock were added aseptically. The mixture was then topped up with sterile distilled water to a final volume of 1 L. This yields **Brock medium** with approximately 0.1% (w/v) N-Z-amine and 0.2% (w/v) dextrin. The medium’s pH was adjusted to 3.5 with sterile 1 M H_2_SO_4_ if necessary. Cultures of *S. acidocaldarius* were grown in this medium under aerobic conditions at 75 °C with shaking (approximately 120 rpm).

### Knock-out of SAMH gene in *H. volcanii*

For the deletion of *hvo_0781*, plasmid pSVA13740 was constructed via *in vivo* ligation using the primers indicated in **Table S3**. Before transformation in *H. volcanii* the obtained plasmid was passed through a dam^-^/dcm^-^ *E. coli* strain for demethylation.^[28]^ Subsequently, *H. volcanii* strain H26 was transformed using polyethylene glycol 600 as described before^[27]^. After colonies were visible on the selective transformation plate a single colony was picked and cultured in 5 mL non-selective Hv-YPC overnight at 45 °C. 5 µL were then used to inoculate a fresh culture in 5 mL Hv-YPC and the culture incubate overnight. This procedure was repeated. From the last culture serial dilutions until 10^-3^ were prepared and 100 µL of each dilution was plated on Hv-Ca-5-FOA agar plates and incubated until visible colonies had grown at 45 °C. Several colonies were transferred on a fresh Hv-YPC plate and incubated for 3 days. Then, half of each of these colonies was used for colony-PCR using the primers indicated in **Table S3**.

### Knock-out of SAMH gene in *S. acidocaldarius*

The deletion plasmid for *saci0719*, pSVA2226, was constructed by a combination of overlap PCR and standard restriction enzyme-based cloning following the manufacturers protocols (NEB) using primers indicated in **Table S3**. Before transformation the integrative deletion plasmid was passed through *E. coli* strain ER1281 for methylation. The methylated plasmid was then transformed into electric competent MW001 cells as described before.^[29]^ Successful transformants were first selected on uracil-free plates and tested by PCR for plasmid integration using primer 3218 and 3219. Positive clones were incubated further in liquid medium and subsequently plated on 5-FOA containing plates. Grown colonies were transferred to uracil containing plates and screened via colony PCR for the deletion of *saci0719*.

### Archaeal culturing and intracellular SAM content determination

*S. acidocaldarius* cultures were grown in Brock media containing uracil (10 µg ⋅ mL). Pre-cultures were inoculated from the cryo-cultures and grown to an OD_600_ of 0.5 (75 °C, 120 rpm, 48 h). Three main cultures were then diluted back in 50 mL Brock to reach an OD_600_ of approximately 0.4-0.5 after 24 h incubation at 75 °C and 120 rpm. Cultures were then divided into two smaller cultures. To one of the two cultures, L-methionine (>98%,Alfa Aesar, Germany) was added to a final concentration of 10 mM, and the cells incubated for another 4 h. Then the end OD_600_ was measured and the cells were harvested at 1900xg for 20 min at 4°C. The supernatant was discarded, and the cell pellet frozen in liquid nitrogen.

*H. volcanii* cultures were grown in CAB medium. Pre-cultures were inoculated with colonies in 5 mL medium and incubated at 45 °C in a rotor overnight. For main cultures the cells were diluted back in 50 ml CAB-medium and treated like the *S. acidocaldarius* cultures with incubation at 45 °C, 120 rpm.

For analysis, the pellets were thawed on ice, re-suspended in 1 mL 10% (*v/v*) perchloric acid and lysed using bead beater Precellys 24 (Bertin Technologies, France) with 2 mL tubes of Precellys Lysing Kit VK05 (Bertin Technologies, France). Bead beating was performed for 3 cycles at 6000 rpm for 20 s with 5 min breaks on ice. The samples were centrifuged for ≥ 15 min at 4 °C and 7227 xg. The supernatant was then prepared for LC-MS/MS analysis.

### Microscopy

To assess the impact of the SAMH deletion on the cell morphology of *H. volcanii* and *S. acidocaldarius* light microscopy was performed.

*Haloferax volcanii* was grown as described before in 20 mL Cab-medium until and OD_600_ of ∼0.05, ∼ 0.3 and ∼1. For microscopy cells were spotted on an 18% saltwater (144 g·L^−1^ NaCl, 18 g·L^−1^ MgCl_2_·6H2O, 21 g·L^−1^ MgSO_4_·7H_2_O, 4.2 g·L^−1^ KCl and 12 mM Tris-HCl, pH 7.5) containing 1% w/v agarose pad. Once the liquid was soaked of by the pad cells were imaged with an in verted microscope (Zeiss Axio Observer.Z1, controlled via Zeiss Blue v.3.3.89) at a 1000 X magnification in phase contrast mode.

For *S. acidocaldarius* 1% agarose was dissolved in Brock medium with a pH of 3.5. Grown cells were spotted on agarose pads and processed like *H. volcanii* with the difference that images were taken in DIC mode.

### Cloning of Overexpression Plasmids

The genes of interest were amplified from genomic DNA via PCR using primers 13185, 13186 for *H. volcanii* and primers 3171, 3172 for *S. acidocaldarius* following the manufacturers protocol (NEB). Successful PCR was confirmed by agarose gel and the PCR product purified from the gel. Subsequently, FX-cloning was performed for the *Hv*SAMH expression construct.^[30]^ The *Saci*SAMH was cloned in the expression vector via restriction enzyme-based cloning. Potentially correct colonies were screened via colony PCR and of positive candidates the plasmid was extracted and sent for sequencing.

### Protein Overproduction

*E. coli* BL21-Gold(DE3) cells were transformed with the plasmids carrying the genes of interest. Pre-cultures were prepared from a single colony and 5 mL LB-medium containing kanamycin (50 μg ⋅ mL^−1^) or ampicillin (100 μg ⋅ mL^−1^), and incubated at 37 °C, 170 rpm overnight. The pre-culture (1%) was then used to inoculate main cultures [400 mL LB-medium containing kanamycin (50 μg ⋅ mL^−1^) or ampicillin (100 μg ⋅ mL^−1)^]. Main cultures were incubated at 37 °C, 170 rpm until an OD_600_ of 0.5–0.7 was reached.

For induction of overexpression, isopropyl-β-D-thiogalactopyranoside (IPTG) was added to a final concentration of 0.4 mM. The overproduction took place at 20 °C, 160 rpm for 20 h. Cells were harvested by centrifugation (4 °C, 7.8×1000 xg, 20 min) and pellets stored at -20 °C until protein purification. Proteins were visualised *via* SDS-PAGE analysis.

### Protein Purification

Pellets were resuspended (4 mL · g^−1^) in lysis buffer [40 mM Tris-HCl, pH 7.4, 500 mM NaCl (*Saci*SAMH) or 1.5 M KCl and 100 mM NaCl (*Hv*SAMH), 10 % (v/*v*) glycerol]. Cell lysis was performed *via* sonication (Branson Sonifier 250, Emerson, St. Louis, MO, USA [duty cycle 50 %, intensity 50 %, 4×30 s with 30 s breaks within]). The lysate was centrifuged for 45 min at 4 °C (24.9 × 1000 xg) to precipitate non-soluble cell fragments. For *Saci*SAMH, the cleared lysate was applied to a nickel-NTA column (resin: HisPur^TM^, Thermoscientific, USA) followed by washing (30 mL lysis buffer containing 10 mM imidazole) and eluting (20 mL lysis buffer with 250 mM imidazole). The protein solution was desalted, using PD-10 columns (GE Healthcare Life Sciences, Little Chalfont, UK) according to the manufacturer’s instructions.

For *Hv*SAMH, the cleared lysate was sterile filtered using Rotilabo^®^ syringe filters (Nylon, 0.45 µm, Carl Roth GmbH, Germany) and applied to a Strep-Tactin Superflow column (IBA Lifesciences GmbH, Germany). The subsequent washing step was performed with 20 mL buffer A, the protein was then eluted using 20 mL buffer A containing 2.5 mM desthiobiotin. Proteins were concentrated with Macrosep Advance Centrifugal Device (30 kDa cut-off, Pall Laboratory, USA).

Protein concentration was determined with a NanoDrop 2000 (Thermo Fisher Scientific, Waltham, MA, USA) at 280 nm. For this, the molecular weight and extinction coefficient (including His_6_-tag) were calculated with the ExPASy ProtParam tool.

### *In vitro* Assay

SAM was purchased from Biosynth. and epimerised by incubation at 75 °C for 2 h. The assays were performed in 50 mM Tris pH 7.5 using 2 mM SAM and 10 µM of the respective SAMH. For the cascade with *Ec*MAT, 50 mM KCl and 20 mM MgCl_2_ were added to the buffer. Selenomethionine and ATP were used in concentrations of 2mM and 3mM, respectively. All enzymes had a concentration of 10 µM. Assays with *Hv*SAMH were performed directly after purification or on the day after with overnight storage at 8 °C. Assays were incubated at 37 °C and 350 rpm for 20 h and stopped by the addition of perchloric acid (final conc. 2.5%). Samples were stored at -20 °C.

### HPLC Analysis

Samples of the *in vitro* assays were first thawed and centrifuged (18213 xg, 4 °C, 45 min) followed by analysis with an Agilent 1100 Series HPLC. An ISAspehr-100-5 C18 AQ column was used with a flowrate of 0.7 mL ⋅ min^-1^. Eluent A was 10 mM NH_4_FA at pH 3.5 and eluent B was acetonitrile. The following gradient method was used: 0-6 min: 100% eluent A, 10-13 min: 80% eluent A, 16-25 min: 100% eluent A. The injection volume was set to 10 µL (method A).

### LC-MS/MS Analysis

Intracellular SAM content was analysed *via* LC-MS/MS. Prior to analysis, samples were diluted 1:10 with ddH_2_O and filtered with cellulose acetate filters (ISERA, Ø 4 mm, pore 0.2 µm). A Sciex 5500+ triplequad LC-MS/MS system was used. Column and eluents were identical to the HPLC method but with a flowrate of 0.4 mL ⋅ min^-1^ and an injection volume of 1 µL. Parameters of the MS method can be found in **Table S5**.

## Acknowledgments

We thank Agnieszka Stanek for initial experiments. This work was funded by the Deutsche Forschungsgemeinschaft (DFG, German Research Foundation) – FOR5596, Project number 510974120. P.N. and S.V.A. were supported by the VW Foundation on a Momentum grant (nr. 94993).

## Author Contributions

A.B., M.K.F.M., P.N., B.W. and L.R. designed and performed experiments. A.B., M.K.F.M., P.N., S.V.A. and J.N.A. wrote the manuscript. The project was conceived by J.N.A, S.V.A. and M.K.F.M. All authors approved the final manuscript.

## References

[1] G. L. Cantoni, ““Biological methylation: selected aspects,”“ Annual review of biochemistry 1975, 44, 435–451.

[2] G. de la Haba, G. A. Jamieson, S. H. Mudd, H. H. Richards, ““S-Adenosylmethionine: The Relation of Configuration at the Sulfonium Center to Enzymatic Reactivity1,”“ J. Am. Chem. Soc. 1959, 81, 3975–3980.

[3] R. T. Borchardt, Y. S. Wu, ““Potential inhibitors of S-adenosylmethionine-dependent methyltransferases. 5. Role of the asymmetric sulfonium pole in the enzymic binding of S-adenosyl-L-methionine,”“ J. Med. Chem. 1976, 19, 1099–1103.

[4] J. R. Matos, C.-H. Wong, ““S-adenosylmethionine: Stability and stabilization,”“ Bioorganic Chemistry 1987, 15, 71–80.

[5] D. F. Iwig, S. J. Booker, ““Insight into the Polar Reactivity of the Onium Chalcogen Analogues of S-Adenosyl-L-methionine,”“ Biochemistry 2004, 43, 13496–13509.

[6] J. L. Hoffman, ““Chromatographic analysis of the chiral and covalent instability of S-adenosyl-L-methionine,”“ Biochemistry 1986, 25, 4444–4449.

[7] G. L. Cantoni, ““ACTIVATION OF METHIONINE FOR TRANSMETHYLATION,”“ Journal of Biological Chemistry 1951, 189, 745– 754.

[8] M. K. F. Mohr, P. Benčić, J. N. Andexer, ““Doping In Vivo Alkylation in E. coli by Introducing the Direct Sulfurylation Pathway of S. cerevisiae,”“ Angew. Chem. Int. Ed. n.d., n/a, e202414598.

[9] A. M. Kunjapur, J. C. Hyun, K. L. J. Prather, ““Deregulation of S-adenosylmethionine biosynthesis and regeneration improves methylation in the E. coli de novo vanillin biosynthesis pathway,”“ Microb Cell Fact 2016, 15, 61.

[10] Y. Lv, J. Chang, W. Zhang, H. Dong, S. Chen, X. Wang, A. Zhao, S. Zhang, Md. A. Alam, S. Wang, C. Du, J. Xu, W. Wang, P. Xu, ““Improving Microbial Cell Factory Performance by Engineering SAM Availability,”“ J. Agric. Food Chem. 2024, 72, 3846– 3871.

[11] M. K. F. Mohr, A. Satanowski, S. N. Lindner, T. J. Erb, J. N. Andexer, ““Rewiring Escherichia coli to transform formate into methyl groups,”“ Microbial Cell Factories 2025, 24, 55.

[12] J. Siegrist, S. Aschwanden, S. Mordhorst, L. Thöny-Meyer, M. Richter, J. N. Andexer, ““Regiocomplementary O-Methylation of Catechols by Using Three-Enzyme Cascades,”“ ChemBioChem 2015, 16, 2576–2579.

[13] H. Deng, D. O’Hagan, ““The fluorinase, the chlorinase and the duf-62 enzymes,”“ Current Opinion in Chemical Biology 2008, 12, 582–592.

[14] T. Kornfuehrer, S. Romanowski, V. de Crécy-Lagard, A. D. Hanson, A. S. Eustáquio, ““An Enzyme Containing the Conserved Domain of Unknown Function DUF62 Acts as a Stereoselective (Rs,Sc)-S-Adenosylmethionine Hydrolase,”“ ChemBioChem 2020, 21, 3495–3499.

[15] A. S. Eustáquio, J. Härle, J. P. Noel, B. S. Moore, ““S-Adenosyl-L-Methionine Hydrolase (Adenosine-Forming), a Conserved Bacterial and Archeal Protein Related to SAM-Dependent Halogenases,”“ ChemBioChem 2008, 9, 2215–2219.

[16] A. S. Eustáquio, F. Pojer, J. P. Noel, B. S. Moore, ““Discovery and characterization of a marine bacterial SAM-dependent chlorinase,”“ Nature Chemical Biology 2008, 4, 69–74.

[17] C. Beaudouin, G. Haurat, J. A. Laffitte, B. Renaud, ““The presence of (+)-S-adenosyl-L-methionine in the rat brain and its lack of effect on phenylethanolamine N-methyltransferase activity.,”“ J Neurochem 1993, 61, 928–935.

[18] C. R. Vinci, S. G. Clarke, ““Homocysteine Methyltransferases Mht1 and Sam4 Prevent the Accumulation of Age-damaged (R,S)-AdoMet in the Yeast Saccharomyces cerevisiae*,”“ Journal of Biological Chemistry 2010, 285, 20526–20531.

[19] L. M. T. Bradbury, M. J. Ziemak, M. El Badawi-Sidhu, O. Fiehn, A. D. Hanson, ““Plant-driven repurposing of the ancient S-adenosylmethionine repair enzyme homocysteine S-methyltransferase,”“ Biochemical Journal 2014, 463, 279–286.

[20] D. Aparici-Carratalá, J. Esclapez, V. Bautista, M.-J. Bonete, M. Camacho, ““Archaea: current and potential biotechnological applications,”“ Research in Microbiology 2023, 174, 104080.

[21] R. U. Haque, F. Paradisi, T. Allers, ““Haloferax volcanii for biotechnology applications: challenges, current state and perspectives.,”“ Appl Microbiol Biotechnol 2020, 104, 1371–1382.

[22] T. D. Brock, K. M. Brock, R. T. Belly, R. L. Weiss, ““Sulfolobus: A new genus of sulfur-oxidizing bacteria living at low pH and high temperature,”“ Archiv für Mikrobiologie 1972, 84, 54– 68.

[23] M. S. Vogt, R. R. Ngouoko Nguepbeu, M. K. F. Mohr, S.-V. Albers, L.-O. Essen, A. Banerjee, ““The archaeal triphosphate tunnel metalloenzyme SaTTM defines structural determinants for the diverse activities in the CYTH protein family,”“ J Biol Chem 2021, 297, 100820.

[24] N. Yu, H. Zhao, W. Wang, M. Dong, ““Enzymatic Fluoroethylation by a Fluoroethyl Selenium Analogue of S-Adenosylmethionine,”“ ACS Catal. 2024, 14, 6211–6216.

[25] I. R. Bothwell, K. Islam, Y. Chen, W. Zheng, G. Blum, H. Deng, M. Luo, ““Se-adenosyl-L-selenomethionine cofactor analogue as a reporter of protein methylation.,”“ J Am Chem Soc 2012, 134, 14905–14912.

[26] R. T. de Silva, M. F. Abdul-Halim, D. A. Pittrich, H. J. Brown, M. Pohlschroder, I. G. Duggin, ““Improved growth and morphological plasticity of Haloferax volcanii.,”“ Microbiology (Reading) 2021, 167,, DOI 10.1099/mic.0.001012.

[27] T. Allers, H.-P. Ngo, M. Mevarech, R. G. Lloyd, ““Development of additional selectable markers for the halophilic archaeon Haloferax volcanii based on the leuB and trpA genes.,”“ Appl Environ Microbiol 2004, 70, 943–953.

[28] J. F. Watson, J. García-Nafría, ““In vivo DNA assembly using common laboratory bacteria: A re-emerging tool to simplify molecular cloning.,”“ J Biol Chem 2019, 294, 15271–15281.

[29] M. Wagner, M. van Wolferen, A. Wagner, K. Lassak, B. H. Meyer, J. Reimann, S.-V. Albers, ““Versatile Genetic Tool Box for the Crenarchaeote Sulfolobus acidocaldarius.,”“ Front Microbiol 2012, 3, 214.

[30] E. R. Geertsma, ““FX cloning: a versatile high-throughput cloning system for characterization of enzyme variants.,”“ Methods Mol Biol 2013, 978, 133–148.

