## Supplementary Information for "(*R*)-*S*-Adenosyl-L-methionine hydrolases counter sulfonium epimerisation in thermophilic archaea"

**Protein sequences**

*SaciS*AMH + **His_6_-tag**

MLKRRTIAILTDYGVSDNYNGTLEGVIRKINPDIDITYITPNAKNFNIFTGAYLLNTSYRYFKKNTIFLVIVDPGVGTQRNAILVKTNNYYFVAPDNGVLYPTIVEDNIDKIIRISNKKFYLSESISNTFHGRDIFAPVASYISVGVDLNVFGEEISKSEIVKLDFSYTLERINENKVKACGKIVYIDHFGNIATTIRNVKLRTGEKALVSLRDKQLELRVVRTFAEGKENELVLYSNGYGFIEIGINMSSASNILGVKEGEDVCIEAYIQEDSNHSTG**SIEGRHHHHHH**

*Hv*SAMH + **Strep-tag**

MITLASDFGLAYPAAMKGVVLSRTDGRLVDVAHDLPRQDVRAAAFWLRETLPYFPPAVHLVVVDPGVGTDRRAVVLRVGDHVFVGPDNGVLRPPARRVAAELDDDAPVEAYEIDVPNPESTTFHGRDVFAPAAADAHVAGASALGSVDRFEPIPVGSLSECRLSEATVSDDGREATGEVLVVDDFGNCITNVPGAFVRDAAGETVWVNGESVPVGRTFEAVARGEKLVTVGSHGNVECDVNHGRGDEAFGLAPGDRVELRVD**ALEWSHPQFEK**

**Table S 1:** Strains used in this study.

| **Strains** | **Background strain** | **Genotype** | **Reference** |
| --- | --- | --- | --- |
| **Archaea** |  |  |  |
| H26 | - | *ΔpyrE2* | ^[1]^ |
| HTQ288 | H26 | *ΔpyrE2 Δhvo_0781* | This study |
| MW001 | - | *ΔpyrEF* | ^[2]^ |
| MW379 | MW001 | *ΔpyrEF Δsaci0719* | This study |
| ***E. coli*** |  |  |  |
| 10-beta  Competent Cells  “TOP10” | - | *Δ(ara-leu) 7697 araD139 fhuA*  *ΔlacX74 galK16 galE15 e14-*  *ϕ80dlacZΔM15 recA1 relA1 endA1*  *nupG rpsL (StrR) rph spoT1 Δ(mrr-*  *hsdRMS-mcrBC)* | New England  Biolabs |
| dam-/dcm-  Competent  Cells | - | *ara-14 leuB6 fhuA31 lacY1 tsx78*  *glnV44 galK2 galT22 mcrA dcm-6*  *hisG4 rfbD1 R(zgb210::Tn10) TetS*  *endA1 rspL136 (StrR) dam13::Tn9*  *(CamR) xylA-5 mtl-1 thi-1 mcrB1*  *hsdR2* | New England  Biolabs |
| ER1821 | - | F^-^ *glnV44 e14^-^(McrA^-^) rfbD1? relA1? endA1 spoT1? Thi-1 Δ(mcrC-mrr)114::IS10* | New England  Biolabs |
| BL21 (DE3) Gold  Competent Cells |  | B F– *omp*T *hsd*S*(*rB– mB–*)* dcm+ Tet^r^ *galλ*(DE3) *end*A Hte | Agilent Technologies, Waldbronn, Germany |

**Table S 2**: Plasmids used in this study.

| **Plasmid** | **Backbone** | **Function** | **Primers used** | **Reference** |
| --- | --- | --- | --- | --- |
| pSVA406 | - | Integrative plasmid for the generation of gene deletions in *S. acidocaldarius* | - | ^[2]^ |
| pSVA2226 | pSVA406 | Knock-out plasmid for the deletion of *saci0719*  (Amp^R^) | 3193, 3194;  3195,  3196 | This study |
| pSA4 | pET15b | Expression vector containing T7 promotor and a 6x His-tag  (Amp^R^) | - | ^[3]^ |
| pSVA2207 | pSA4 | Plasmid for the expression of Saci0719-6xHis in *E. coli*  (Amp^R^) |  | This study |
| pTA131 | - | Integrative plasmid for the generation of gene deletions in *H. volcanii* | - | ^[4]^ |
| pSVA13740 | pTA131 | Knock-out plasmid for the deletion of *hvo_0781*  (Amp^R^) | 13177, 13178; 13179, 13180 | This study |
| p7XC3S | - | FX cloning  vector with a T7 promoter and C-  terminal 3C protease cleavage  site and Strep-tag  (Kan^R^) | - | ^[5]^ |
| pSVA13786 | p7XC3S | Plasmid for the expression of HVO_0781-Strep in *E. coli*  (Kan^R^) | 13185, 13186 | This study |

**Table S 3:** Primers used in this study.

| **Primer** | **Sequence** | **Description** |
| --- | --- | --- |
| 11066 | TCTAGAGCGGCCGCCAC | Forward primer for linearization of pTA131 |
| 11067 | GGTACCCAATTCGCCCTATAGTG | Reverse primer for linearization of pTA131 |
| 13177 | GGCGAATTGGGTACCACCACCGAATGGTCGAAGAG | Forward primer for the amplification of ~500 bp  up-stream region of *hvo_0781* with 15 complementary bases to  linearized pTA131 |
| 13178 | AAGCAGCGTCGCCATCTCAACAGGGGTTGGGGTC | Reverse primer for the amplification of ~500 bp up-stream region of *hvo_0781* with 15 complementary bases to the *hvo_0781* down-stream region |
| 13179 | CCAACCCCTGTTGAGATGGCGACGCTGCTTTTC | Forward primer for the amplification of ~500 bp  down-stream region of *hvo_0781* with 15 complementary bases to  the up-stream region |
| 13180 | GGCGGCCGCTCTAGAGGCGTCGCGTAGAAGTCG | Reverse primer for the amplification of ~500 bp  down-stream region of *hvo_0781* with 15 complementary bases to  linearized pTA131 |
| 13185 | atatatGCTCTTCtAGTattacgctcgcgtccgacttcgggctc | Forward primer for the amplification of *hvo_0781* with a SapI restriction site |
| 13186 | tatataGCTCTTCaTGCAtcAacAcggagttcgacgcggtc | Reverse primer for the amplification of *hvo_0781* with a SapI restriction site |
| 13197 | CGCGTTTCGACCTTCCTACC | Forward primer for the screening of *hvo_0781*  deletion |
| 13198 | AGCACCCGAACCGAGAAGAC | Reverse primer for the screening of *hvo_0781*  deletion |
| 3193 | GGTCCATGGGCCATTGGACCTTCTCCAGCT | Forward primer for the amplification of ~500 bp  up-stream region of *saci0719* with an NcoI restriction site |
| 3194 | CTTCACCTTCTTTTACTCCCTCTAAGGTTCCATTGTAATTATC | Reverse primer for the amplification of ~500 bp  up-stream region of *saci0719* with ~15 complementary bases to  the down-stream region |
| 3195 | CAATGGAACCTTAGAGGGAGTAAAAGAAGGTGAAGATGTTTGC | Forward primer for the amplification of ~500 bp  down-stream region of *saci0719* with ~15 complementary bases to  the upstream region |
| 3196 | GGTGGATCCGCATTGATGAAGCATTTTCCTCTG | Reverse primer for the amplification of ~500 bp  down-stream region of *saci0719* with an BamHI restriction site |
| 3218 | GTTGCACCCAAGCTAGG | Forward primer for the screening of *saci0719*  deletion |
| 3219 | GATGCCAAACGTTGCTAG | Reverse primer for the screening of *saci0719*  deletion |
| 3171 | GGACCATGGCCCTAAAAAGACGAACAATAGCAATAC | Forward primer for the amplification of *saci0719* with an NcoI restriction site |
| 3172 | GGCGGATCCTGTGGAATGGTTGGAATCTTCCTG | Reverse primer for the amplification of *saci0719* with an BamHI restriction site |

**HPLC analysis**

**Table S 4:** Retention times in HPLC method A as identified by authentic standards.

| Molecule | Retention time / min |
| --- | --- |
| Adenine | 13.5 |
| Adenosine | 14.6 |
| AMP | 13.3 |
| ADP | 6.8 |
| ATP | 5.2 |
| 5´-Chloro-5´-deoxyadenosine | 16.2 |
| Inosine | 14.0 |
| 5´-Methylthioadenosine | 16.7 |
| (*S*_S_,*S*_Cα_)-SAM | 8.5 |
| (*R*_S_,*S*_Cα_)-SAM | 9.0 |

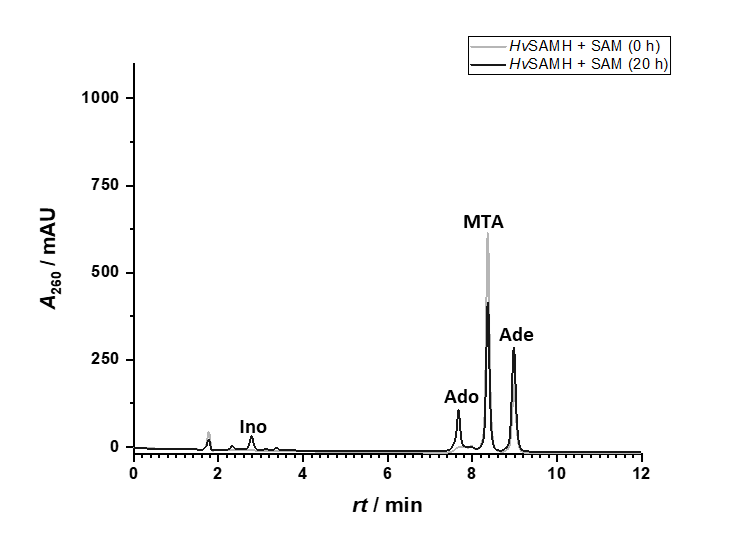
To confirm the formation of inosine, the HPLC method of Köppl et al. was used (method B).^[6]^

**Figure S 1:** HPLC chromatogram (method B) showing the production of adenosine and inosine.

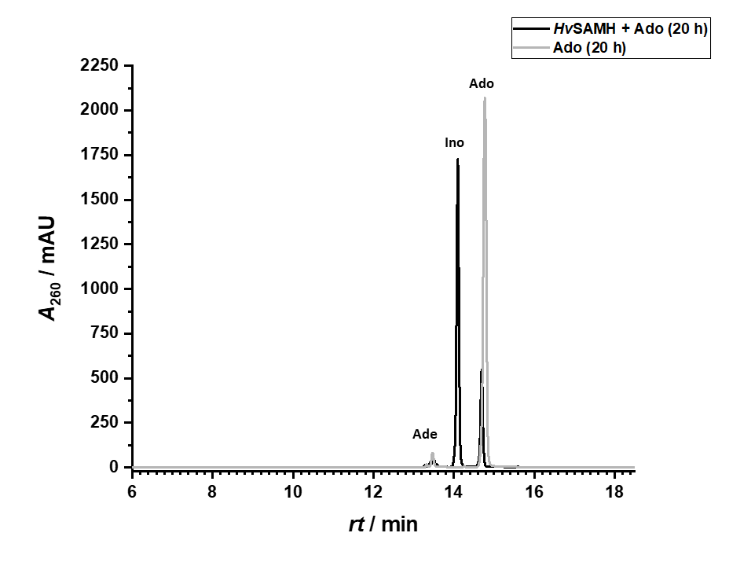

**Figure S 2:** Confirmation of deaminase side reactivity. HPLC chromatogram (method A) of HvSAMH and Ado after 20 h of incubation at 37 °C.

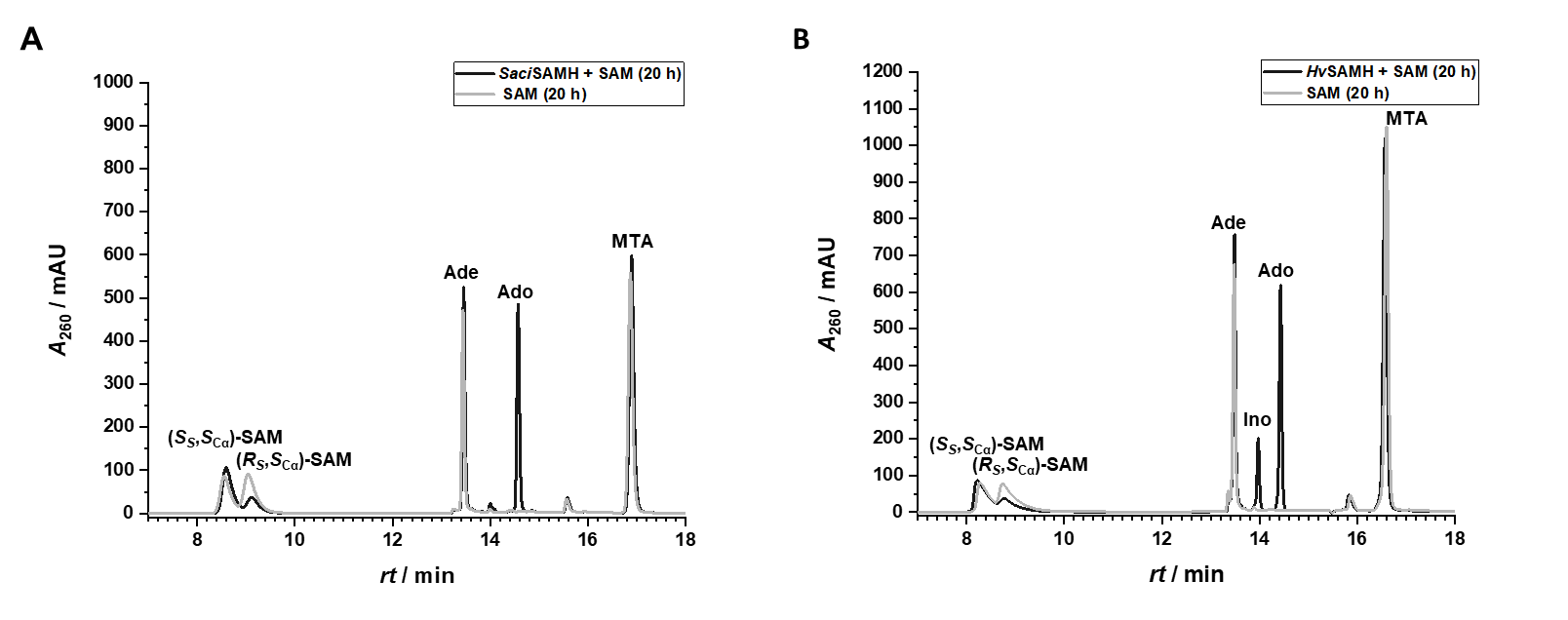

**Figure S 3:** Confirmation of no chlorinase activity of SaciSAMH (A) and HvSAMH (B). HPLC chromatograms (method A) of respective SAMH and SAM after 20 h of incubation at 37 °C (different section of chromatograms shown in Figure 2).

**LC-MS analysis**

The LC-MS/MS method is based on the method by Zhang et al.^[7]^

**Table S 5:** Setting of the LC-MS/MS method to analyse intracellular SAM content. CE = collision energy, CXP = cell exit potential.

| **MS settings** | Curtain gas | 20 psi |
| --- | --- | --- |
|  | Ion spray voltage | +5500 V |
|  | Source temperature | 500 °C |
|  | Ion source gas 1 | 40 psi |
|  | Ion source gas 2 | 50 psi |
|  | Declustering potential | 89.2 V |
|  | Entrance Potential | 14.0 V |
|  | Observed mass transitions (specific for SAM) | *m/z* 399.15 → *m/z* 135.9,  CE: 32.0 V, CXP: 47.0 V |
|  |  | *m/z* 399.15 → *m/z* 250.02,  CE: 23.0 V, CXP: 47.9 |
|  |  | *m/z* 400.15 → *m/z* 250.02,  CE: 23 V, CXP: 47.9 V |
| **LC settings** | Sample volume injected | 1 µL |
|  | Equilibration | 100% A |
|  | Gradient | 18 min 100% A |
|  |  | 1 min to 100% B |
|  |  | 6 min 100% B |
|  |  | 11 min to 100% A |
|  |  | 14 min 100% A |

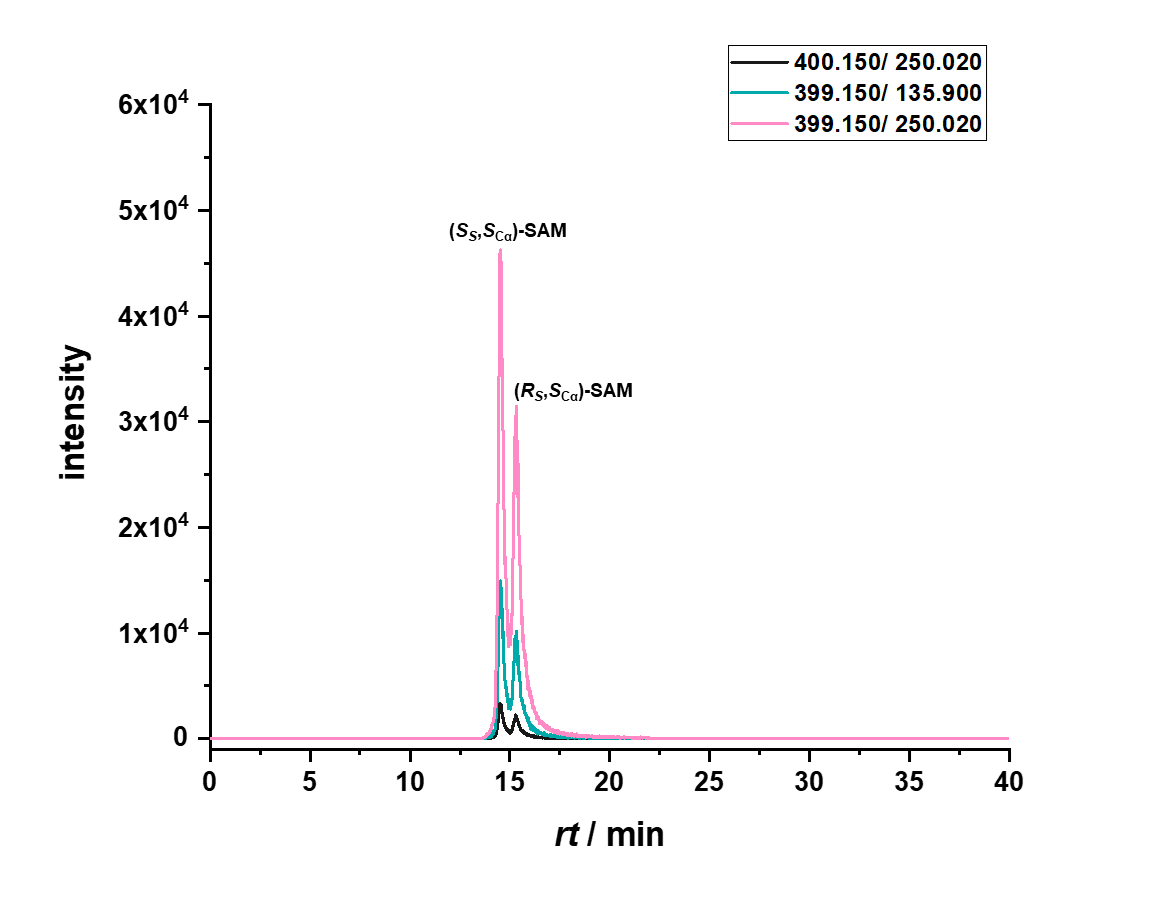

**Figure S 4:** MRM-Scan of epimerised SAM standard (10 µm). m/z of daughter ions and fragments are given in the legend.

**SDS-PAGE analysis of protein purification**

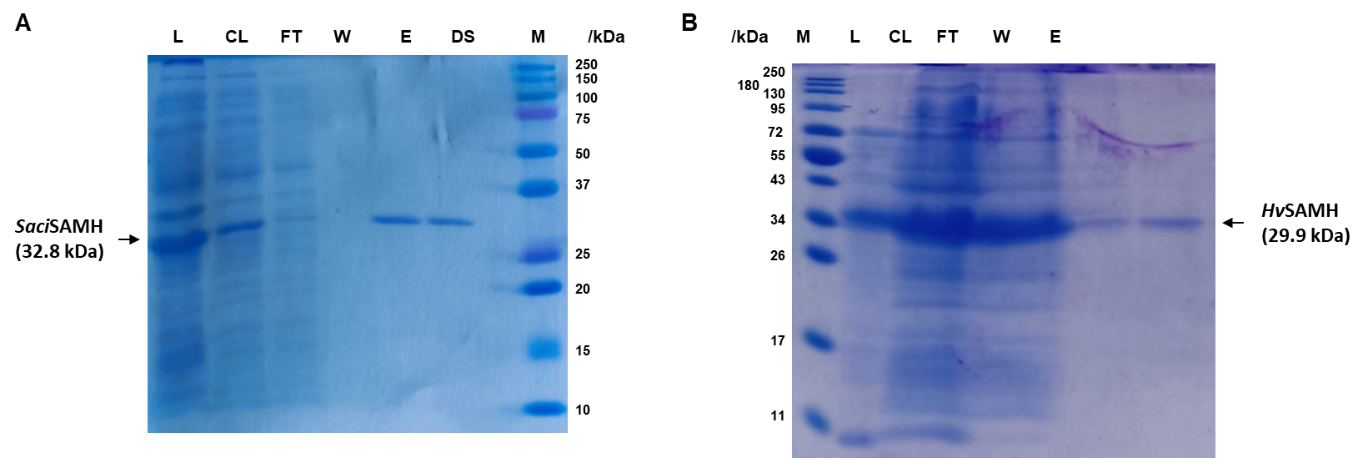

**Figure S 5:** SDS-PAGE analysis of protein purification of SaciSAMH (A) and HvSAMH (B). L: Lysate, CL: Cleared lysate, FT: Flowthrough, W: Wash, E: Eluate, DS: Desalted protein, M: marker [Precision Plus Protein^TM^ Standards Dual Color, BIORAD (A), Blue Prestained Protein Standard, Broad Range, New England BioLabs (B).

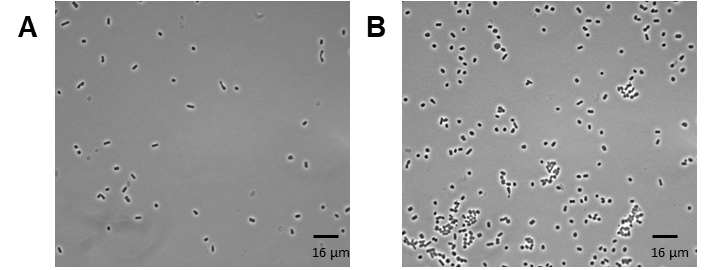

**Figure S 6**: Phase contrast microscopy images of Hv_WT and Hv_KO. Cells were grown in shaking culture at 45 °C to an OD_600_ = 0.25 (Hv_WT) and OD600 = 0.35 (Hv_KO). The scale bar indicates 16 µm.

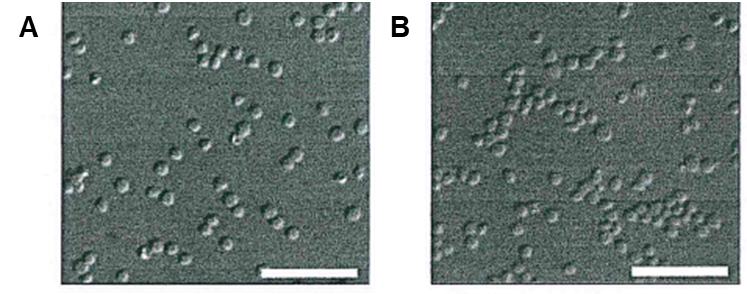

**Figure S 7:** DIC III light microscopy images of Saci_WT (A) and Saci_KO (B). Cells were grown in a shaking culture at 75 °C to an OD_600_  of ~0.3. The scale bar indicates 10 µm.

**References**

[1]	G. Bitan-Banin, R. Ortenberg, M. Mevarech, ““Development of a gene knockout system for the halophilic archaeon Haloferax volcanii by use of the pyrE gene.,”” *J Bacteriol* **2003**, *185*, 772–778.

[2]	M. Wagner, M. van Wolferen, A. Wagner, K. Lassak, B. H. Meyer, J. Reimann, S.-V. Albers, ““Versatile Genetic Tool Box for the Crenarchaeote Sulfolobus acidocaldarius.,”” *Front Microbiol* **2012**, *3*, 214.

[3]	S.-V. Albers, Z. Szabó, A. J. M. Driessen, ““Archaeal homolog of bacterial type IV prepilin signal peptidases with broad substrate specificity.,”” *J Bacteriol* **2003**, *185*, 3918–3925.

[4]	T. Allers, H.-P. Ngo, M. Mevarech, R. G. Lloyd, ““Development of additional selectable markers for the halophilic archaeon Haloferax volcanii based on the leuB and trpA genes.,”” *Appl Environ Microbiol* **2004**, *70*, 943–953.

[5]	E. R. Geertsma, ““FX cloning: a versatile high-throughput cloning system for characterization of enzyme variants.,”” *Methods Mol Biol* **2013**, *978*, 133–148.

[6]	L.-H. Koeppl, D. Popadić, R. Saleem-Batcha, P. Germer, J. N. Andexer, ““Structure, function and substrate preferences of archaeal S-adenosyl-l-homocysteine hydrolases,”” *Communications Biology* **2024**, *7*, 380.

[7]	J. Zhang, J. P. Klinman, ““High-performance liquid chromatography separation of the (S,S)- and (R,S)-forms of S-adenosyl-l-methionine,”” *Analytical Biochemistry* **2015**, *476*, 81–83.
